## Supplementary materials for "Mothers front-load their investment to the egg stage when helped in a wild cooperative bird"

**Supplementary materials include: (details in next page)**

Supplementary text
Figures S1 to S7
Tables S1 to S8
References

**Supplementary materials include:**

#### A - Identification of the time windows of effect for temperature and rainfall

*Rationale and methodology for the sliding window approach*

To control for the effects of variation in environmental temperature and rainfall on egg volume and maternal provisioning rate in the models presented in the main paper, we fitted two predictors within each model: a “heat waves” index (the total number of days within a specific time window prior to the focal event [laying or provisioning] in which the maximum daily temperature exceeded 35°C) and a rainfall index (the total amount of rainfall that fell within a specific time window prior to the focal event). The ‘heat waves’ index as defined here (i.e., number of days above 35°C) has been shown to appropriately capture hot-weather events in the Kalahari and it impacts the reproductive biology of several Kalahari bird species [1,2]. As the timing and duration of these specific time windows of effect were not known and could well differ between the two indices (heat waves and rainfall) and across the two response terms (egg volume and provisioning rate), prior to proceeding with the modelling exercise described in the main paper we used a sliding window approach to objectively identify these windows [3]. The sliding window approach described below was applied four times, to identify the best-supported time window of effect for each of the two indices when modelling each of the two responses. During this process we allowed for a linear effect of the heat wave index and both linear and quadratic effects of the rainfall index.

Within each application of the sliding window approach, we considered all possible temporal windows of >4 days in length, between an earliest start date of 80 days prior to egg laying and a latest end date of the day of egg laying. Only sliding windows of >4 days in length were considered, in order to decrease the likelihood of false positive results (which are more probable for very short windows). For each possible time window, the focal environmental index (heat waves or rainfall; see above) was calculated for all breeding attempts in the data set and then fitted as an additional predictor in a ‘baseline model’ for the focal response term (which was the full model described in the main paper for that response term, but lacking the heat waves and rainfall predictors). The level of statistical support for this time window was then calculated as the ΔAIC value between this model and the baseline model without this new predictor (‘AIC support’ below). The level of AIC support for all of the different possible windows for the focal environmental index were then ranked and the best-supported window carried forward for use within the model described in the main paper if it improved the fit of the baseline model by >6 AIC points (a conservative threshold to avoid the accidental inclusion of uninformative terms [4]).

To assess the likelihood that this sliding window approach had yielded a false positive result for the best-supported window in each case, we carried out 25 randomisations of the data set [3]. In each randomisation, the 'biological reference date' (in this case the lay date of the focal clutch) in the data set was randomised by re-shuffling, similar to the approach implemented in the R package 'climwin' [5]. Following each randomisation of the data set, the full sliding window protocol described above was applied for the focal environmental variable, and the 'AIC support' for the model containing the best-supported window of effect for that environmental variable was recorded. The likelihood that the sliding window identified using the real (non-randomised) data set arose by chance (i.e., as a false positive) could then be estimated by calculating the proportion of the randomisations that yielded a best-supported window with stronger AIC support than that identified using the real data set.

*The time windows of effect identified for the heat wave and rainfall effects on egg volume*

The sliding window analysis for the effect of heat waves on egg volume identified a best-supported window that spanned the 13 days prior to egg laying (AIC support = -11.13), for which the heat wave index had a negative effect on egg volume (Figure S2a). The analysis identified a single and localised AIC peak in the sliding window landscape (Figure S2b) with no clear alternative windows supported by the data. None of our 25 randomisations yielded a best-supported window with equal or stronger AIC support than that identified using the real data, indicating that this result is unlikely to have arisen by chance (Figure S2c). Inclusion of this heat waves index as a predictor in the full model presented in the main paper led to a significant negative effect on egg volume (Table 1). High temperatures have been shown to negatively predict egg size in other species too [6–8]. It is conceivable that this pattern reflects an adaptive strategy (e.g., smaller eggs may be easier to keep cool) but could also reflect a detrimental effect of heat stress on pre-laying maternal condition or physiology [7,9].

The sliding window analysis for the effect of total rainfall on egg volume identified a best-supported window that spanned 44 to 49 days prior to egg laying (AIC support = -12.90), for which the rainfall index had a quadratic effect on egg volume (Figure S3a). This analysis revealed more scattered AIC support across the sliding window landscape without such a clear single peak (Figure S3b), and three of our 25 randomisations yielded a best-supported window of effect with equal or stronger AIC support than that identified using the real data, indicating a 12% probability that this reflects a false positive result (Figure S3c). Inclusion of this total rainfall index as a predictor in the full model presented in the main paper led to a significant negative quadratic relationship between rainfall index and egg volume (Tables 1). The shape of the quadratic rainfall relationship detected (Figure S3a) is consistent with the general expectation of a beneficial effect of rainfall on resource availability in this arid environment (e.g., see the positive effect of rainfall on clutching rate per year, Tables S7 & S8 below), but costs associated with very high rainfall events (that can damage the birds' woven structures and flood the landscape).

Additional analyses confirmed that the female helper number effects on egg volume detected in the main text have not been determined by including the sparser data for higher rainfall values or the resulting quadratic rainfall fit. When we just fit a linear rainfall relationship to the egg volume data from below the peak in the detected quadratic rainfall relationship (i.e., for rainfall levels < 60mm; see Figure S3a), where the data density is higher, the effect sizes for the effects of both female helper number (i.e., prior to partitioning) and Δ female helper number (following partitioning) on egg volume are virtually unchanged: (i) Female helper number effect size ± SE with full data set and quadratic rainfall fit (as per Table 1 in the manuscript) = 0.018 ± 0.009 $cm^3$ / female helper, and then for rainfall < 60mm and linear rainfall fit = 0.019 ± 0.009 $cm^3$ / female helper (the associated p-value for female helper number drops slightly from 0.038 to 0.026). (ii) Δ female helper number effect size ± SE with full data set and quadratic rainfall fit (as per Table 2) = 0.019 ± 0.009; and then for rainfall < 60mm and linear rainfall fit = 0.019 ± 0.009 $cm^3$ / female helper (the associated p-value for female helper number drops slightly from 0.037 to 0.035). Not including rainfall effects in the egg volume models leaves the effect size estimates for both female helper number (i.e., prior to partitioning) and Δ female helper number (following partitioning) virtually unchanged: Female helper number effect size ± SE with rainfall index included (as per Table 1) = 0.018 ± 0.009 cm3 / female helper, and without rainfall included = 0.017 ± 0.009 cm3 / female helper (the associated p-value rises slightly from 0.038 to 0.063). Δ female helper number effect size ± SE with rainfall included (as per Table 2) = 0.019 ± 0.0090 cm3 / female helper, and without rainfall included = 0.017 ± 0.0093 cm3 / female helper (the associated p-value rises slightly from 0.037 to 0.063).

*The time windows of effect identified for the heat wave and rainfall effects on maternal provisioning rate*

The sliding window analysis for the effect of heat waves on maternal provisioning rate identified a best-supported window that spanned 51-59 days prior to egg laying (AIC support = -10.01), for which the heat wave index had a positive effect on maternal provisioning rate (Figure S4a). The analysis revealed AIC support that was widely distributed across the sliding window landscape (Figure S4b), but none of our 25 randomisations yielded a best-supported window of effect with equal or stronger AIC support than that identified using the real data (Figure S4c). Inclusion of this heat waves index as a predictor in the full model presented in the main paper led to a significant positvie effect on maternal provisioning rate (Tables 3). Our inferences regarding the effects of female and male helper number on maternal provisioning rate remain unchanged, however, if this heat waves predictor is excluded from the model in the main paper.

The sliding window analysis for the effect of total rainfall on maternal provisioning rate identified a best-supported window that spanned 61-78 days prior to egg laying (AIC support = -13.08), for which total rainfall had a positive quadratic effect on maternal provisioning rate (Figure S5a). The analysis revealed a clear peak of AIC support in the sliding window landscape (Figure S5b). Four of our 25 randomisations yielded a best-supported window of effect with equal or stronger AIC support than that identified using the real data, indicating a 16% probability that this reflects a false positive result (Figure S5c). Inclusion of this total rainfall index as a predictor in the full model presented in the main paper led to a significant positive quadratic relationship between rainfall index and maternal provisioning rate (Tables 3). The quadratic rainfall relationship detected (Figure S5a) is suggestive of an accelerating positive effect of increasing rainfall on maternal provisioning rate, consistent again with beneficial effects of rainfall on resource availability in this arid environment. Not including rainfall effects in models for maternal provisioning rates leaves the effect size estimates for both female helper number and Δ female helper number virtually unchanged: Female helper number effect size with rainfall included = -0.457 ± 0.195 feeds / hour / female helper (as per Table 3), and without rainfall included = -0.454 ± 0.202 feeds / hour /

female helper (the associated p-value rises slightly from 0.020 to 0.026). Δ female helper number effect size with rainfall included = -0.559 ± 0.269 feeds / hour / female helper (as per Table 4), and without rainfall = -0.530 ± 0.279 feeds / hour / female helper (the associated p-value rises slightly from 0.040 to 0.060).

**B – No evidence of maternal adjustment of number of clutches laid per year according to helper numbers**

We found no evidence that mothers adjusted the total number of clutches that they laid per breeding season according to the average number of female or male helpers that they had in their group over the course of the breeding season. Analysis at the population level revealed that the number of clutches that a mother laid per breeding season was not significantly predicted by either the number of female helpers (female helper number effect ± SE = 0.002 ± 0.038 clutches / female helper, $\chi^2_1 < 0.01$, $p = 0.963$) or the number of male helpers (male helper number effect ± SE = 0.042 ± 0.045 clutches / male helper, $\chi^2_1 = 0.86$, $p = 0.355$). We found similar results after partitioning variation in helper numbers into their within- and among-mother components; the number of clutches laid was not significantly predicted by within-mother variation in either female helper number (Δ female helper number effect ± SE = -0.055 ± 0.049 clutches / female helper; $\chi^2_1 = 1.26$, $p = 0.262$) or male helper number (Δ male helper number effect ± SE = -0.039 ± 0.055 clutches / male helper; $\chi^2_1 = 0.50$, $p = 0.479$). The number of clutches that a mother laid per breeding season was significantly positively predicted by the total rainfall that fell during that breeding season (effect size ± SE = 0.001 ± 0.0003 clutches / mm of rainfall; $\chi^2_1 = 11.51$, $p < 0.001$).

**C – Maternal plasticity in egg volume cannot be readily attributed to carry-over effects of past help**

The apparent maternal plasticity in egg size according to female helper number detected in the main paper could conceivably arise not because mothers *pre-emptively* adjust egg investment to the expected level of post-natal helping that a clutch will receive, but because the *past* actions of helpers in a previous breeding attempt have impacted maternal condition at laying (e.g., via lightening maternal workloads [10–12]). This alternative scenario cannot readily explain our findings, however, as (i) maternal body condition before laying is not predicted by the helper numbers that she had during her previous breeding attempt (see Supplementary materials D, below); (ii) the time since the last breeding attempt does not predict egg volume (either in isolation or via interactions with current helper numbers), suggesting that egg volume is not appreciably impacted by carry-over effects of past reproductive effort (see Supplementary materials E, below); and, the number of helpers in the previous breeding attempt does not predict egg volume (either in isolation or via interactions with the time since the last breeding attempt) (see Supplementary materials F, below). As such, it seems more likely that the egg size plasticity observed does reflect maternal adjustment of pre-natal investment according to the likely future availability of post-natal help [13,14], which is highly predictable at the time of laying (Figure S1).

**D – No evidence of carry-over effects of past help on maternal body condition at laying**

We investigated whether maternal body condition before laying a given clutch was predicted by the numbers of helpers present during the rearing of her previous clutch. We restricted the analysis to those 'previous' clutches in which at least one nestling fledged, in order to ensure that mothers had had to engage in post-natal care during the previous breeding attempt and that helpers had had the opportunity to lighten the maternal post-natal workload. We routinely measured body mass (g) and tarsus length (mm) of mothers caught throughout the study period. We used mass measurements from mothers up to 45 days before they produced a new focal clutch (i.e., pre-laying). The data set comprised 45 focal clutches laid by 33 mothers in 26 social groups. We calculated the scale mass index of body condition (hereafter 'maternal pre-laying body condition') following [15], and built a linear mixed model to explain variation in this variable. We investigated the effects of female and male helper number in the previous breeding attempt (as fixed effects) on maternal pre-laying body condition. We also included a fixed effect for the number of previous clutches laid by the mother during that breeding season. Mother ID, social group ID and breeding season ID were included as random effect intercepts. We found no evidence that maternal pre-laying body condition is predicted by the number of female helpers (effect of female helper number on maternal pre-laying condition ± SE = -0.23 ± 0.35 g / female helper, $\chi^2_1 = 0.36$, p = 0.551) or male helpers (effect of male helper number on maternal pre-laying condition ± SE = 0.11 ± 0.40 g / male helper, $\chi^2_1 = 0.03$, p = 0.862) that she had in her previous breeding attempt.

**E – Female helper effects on egg volume do not depend on time since the last breeding attempt or the number of helpers**

If mothers laid larger eggs when assisted by more female helpers (the relationship observed in the main paper) because female helper contributions lightened the mothers' post-natal workload within the *previous* breeding attempt, we would expect the positive effect of female helper number on egg volume to decrease in magnitude with increasing time since the last breeding attempt (i.e., an interaction between female helper number and time since last breeding attempt within the egg volume model). To investigate whether this was the case, we re-fitted the egg volume model presented in the main text to include 'time since last breeding attempt' (range 36-408 days; mean = 86.68 days) as an additional fixed effect predictor. We restricted the data set to only include the egg volumes of those breeding attempts for which the previous breeding attempt had fledged at least one nestling, in order to ensure that mothers had had to engage in post-natal care during the previous breeding attempt and that helpers had had the opportunity to lighten the maternal post-natal workload (n = 136 eggs from 79 clutches laid by 40 mothers in 32 groups). Time since the last breeding attempt did not explain variation in egg volume, either as a simple predictor (effect size ± SE = 0.22 ± 0.34 $cm^3$ / day elapsed, $\chi^2_1 = 0.37$, p = 0.544) or as an interaction with female ($\chi^2_1 = 0.01$, p = 0.909) or male helper number ($\chi^2_1 = 0.02$, p = 0.875).

**F – Variation in egg volume is better explained by 'current' number of helpers than by number of helpers in the previous breeding attempt**

If mothers laid larger eggs when assisted by more female helpers (the relationship observed in the main paper) because female helper contributions lightened the mothers' post-natal workload within the *previous* breeding attempt, we would expect that the number of helpers in the previous breeding attempt explains variation in egg volume better than the number of helpers in the current breeding event. To test this hypothesis, we used AIC to compare (a) the egg volume model presented in the main text including 'time since last breeding' as an additional fixed effect predictor in isolation and in interaction with 'current breeding attempt' male and female helper number (this is the model presented in Section E, above), and (b) an identical model replacing 'current breeding attempt' male and female helper numbers by 'previous breeding attempt' male and female helper numbers. Both, previous and current helper number variables were not included in the same model due to their high correlation (Pearson's product-moment correlation [95% CI] = 0.79 [0.71,

0.85] for previous and current helper number, for both male and female helpers). However, if helper effects on egg volume were a carry-over effect of past help, we would expect the 'previous breeding attempt' model to outperform (i.e., have lower AIC) the 'current breeding attempt' model. AIC model comparison provides an ideal framework for this test as it allows statistical comparison of non-nested models. Again, we restricted the dataset to only include the egg volumes of those breeding attempts for which the previous breeding attempt had fledged at least one nestling, in order to ensure that mothers had had to engage in post-natal care during the previous breeding attempt and that helpers had had the opportunity to lighten the maternal post-natal workload (n = 113 eggs from 66 clutches laid by 38 mothers in 32 groups with information for current and previous breeding attempt helper numbers). The 'previous breeding attempt' model (AIC = 1575.35) did not outperform the 'current breeding attempt' model (AIC = 1574.87). In fact, it fitted the data slightly worse than the 'current breeding attempt' model providing no evidence that helper effects on egg volume could be mediated by carry-over effects of past help.

#### Supplementary Figures

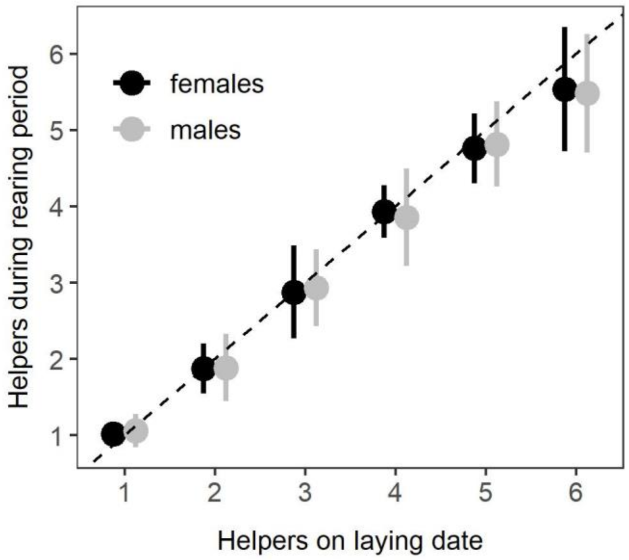

**Figure S1. Number of helpers at laying predicts the number of helpers during the nestling rearing period.** Mean ± standard deviation (SD) is presented for both male and female helper numbers (dashed line indicates a 1:1 relationship). For female helper number, Linear model: N = 271 breeding attempts, β = 0.94 ± 0.017. For male helper number, Linear model: N = 271 breeding attempts, β = 0.93 ± 0.022).

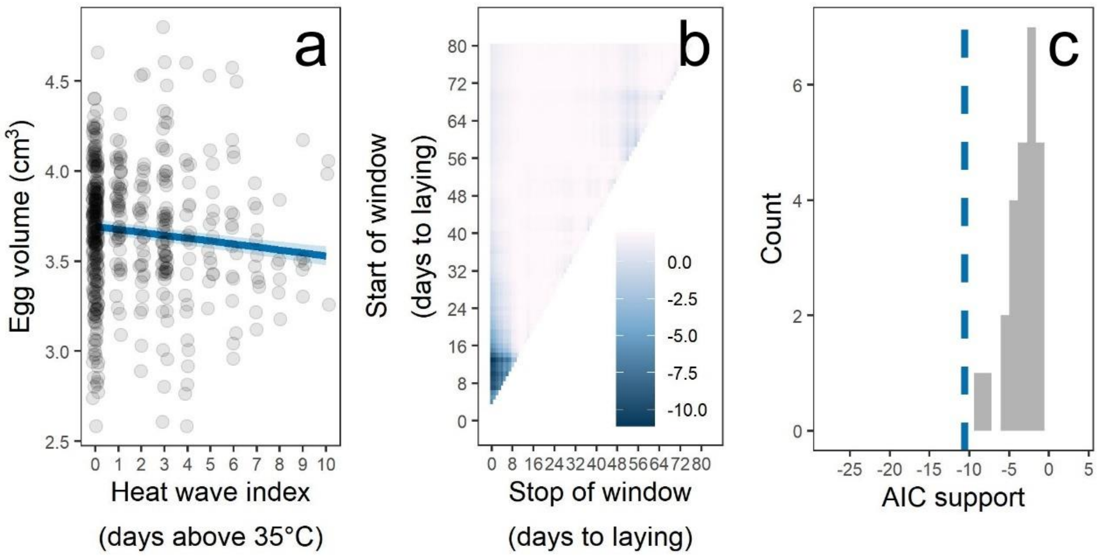

**Figure S2. Sliding window analysis for the effect 'heat waves' (days above 35°C) on egg volume**. See Supplementary materials A above for methods and interpretation. (a) Effect of the best-supported 'heat waves' index (i.e., that calculated for 0-13 days prior to egg laying) on egg volume when tested within the baseline model. Raw data points in black and regression line (± SE) in blue. (b) AIC support (i.e., the difference in AIC between a given sliding window model and the baseline model) for all possible sliding windows of >4 days in length within the 80 days before egg laying. The darker the colour of the tiles, the stronger the support for a given window. (c) Histogram showing the AIC support for the best-supported heat wave index windows arising from 25 randomisations (i.e., the distribution of AIC support expected if no relationship exists between the heat waves index and egg volume). The blue dashed line illustrates the AIC support achieved using the best-supported window from the real data set.

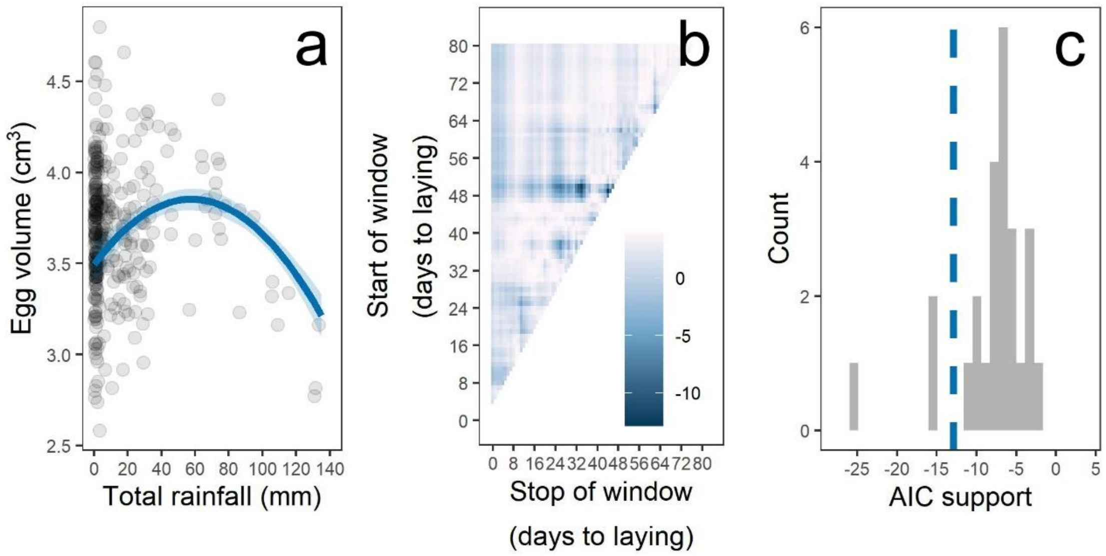

**Figure S3. Sliding window analysis for the effect of total rainfall (mm) on egg volume**. See Supplementary materials A above for methods and interpretation. (a) Effect of the best-supported total rainfall index (i.e., that calculated for 49-44 days prior to egg laying) on egg volume when tested within the baseline model. Raw data points in black and regression line (± SE) in blue. (b) AIC support (i.e., difference in AIC between a given sliding window model and the baseline model) for all possible sliding windows of >4 days in length within the 80 days before egg laying. The darker the colour of the tiles, the stronger the support for a given window. (c) Histogram showing the AIC support for the best-supported rainfall index windows arising from the 25 randomisations (i.e., the distribution of AIC support expected if no relationship exists between the rainfall index and egg volume). The blue dashed line illustrates the AIC support achieved using the best-supported window from the real data set.

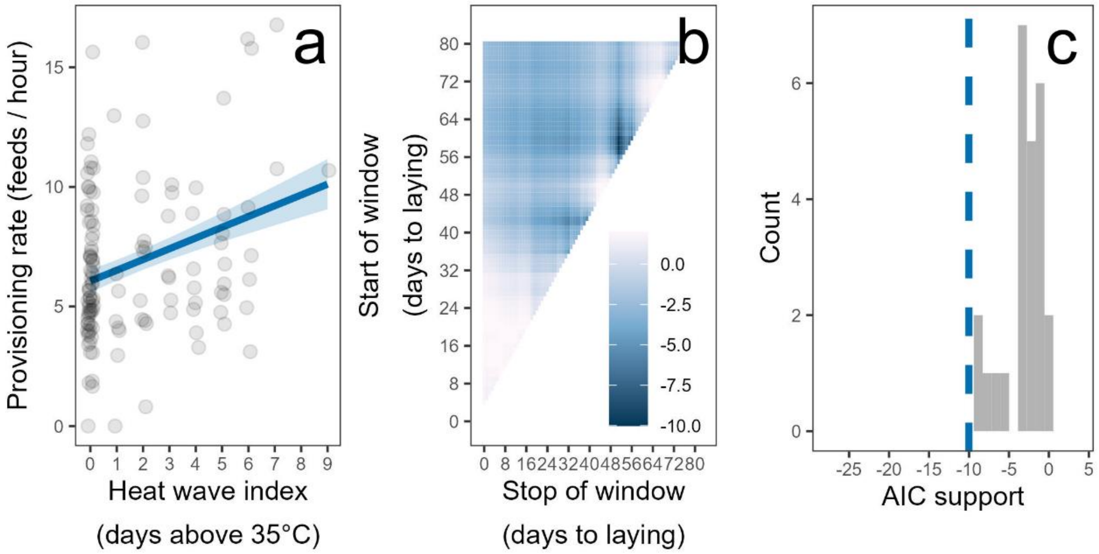

**Figure S4**. **Sliding window analysis for the effect of 'heat waves' (days above 35°C) on maternal provisioning rate**. See Supplementary materials A above for methods and interpretation. (a) Effect of the best-supported 'heat waves' index (i.e., that calculated for 59-51 days prior to egg laying) on maternal provisioning rate when tested within the baseline model. Raw data points in black and regression line (± SE) in blue. (b) AIC support (i.e., difference in AIC between a given sliding window model and the baseline model) for all possible sliding windows of >4 days in length within the 80 days before egg laying. The darker the colour of the tiles, the stronger the support for a given window. (c) Histogram showing the AIC support for the best-supported heat waves index windows arising from 25 randomisations (i.e., the distribution of AIC support expected if no relationship exists between the heat waves index and maternal provisioning rate). The blue dashed line illustrates the AIC support achieved using the best-supported window from the real data set.

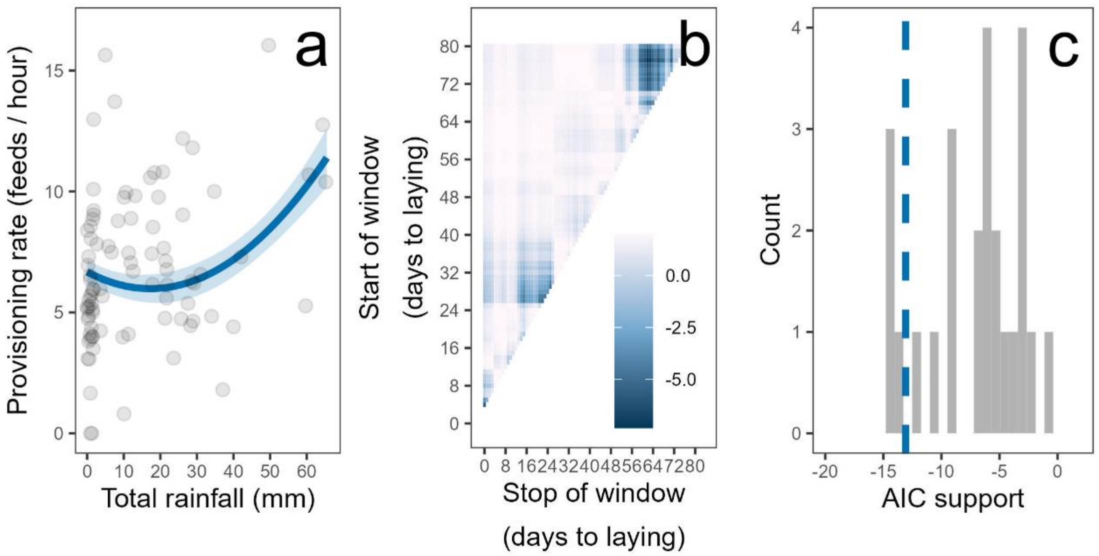

**Figure S5. Sliding window analysis for the effect of total rainfall (mm) on maternal provisioning rate**. See Supplementary materials A above for methods and interpretation. (a) Effect of the best-supported total rainfall index (i.e., that calculated for 78-61 days prior to egg laying) on maternal provisioning rate when tested within the baseline model. Raw data points in black and regression line (± SE) in blue. (b) AIC support (i.e., difference in AIC between a given sliding window model and the baseline model) for all possible sliding windows of >4 days in length within the 80 days before egg laying. The darker the colour of the tiles, the stronger the support for a given window. (c) Histogram showing the AIC support for the best-supported rainfall index windows from each of the 25 randomisations (i.e., the distribution of AIC support expected if no relationship exists between the rainfall index and maternal provisioning rate). The blue dashed line illustrates the AIC support achieved using the best-supported window from the real data set.

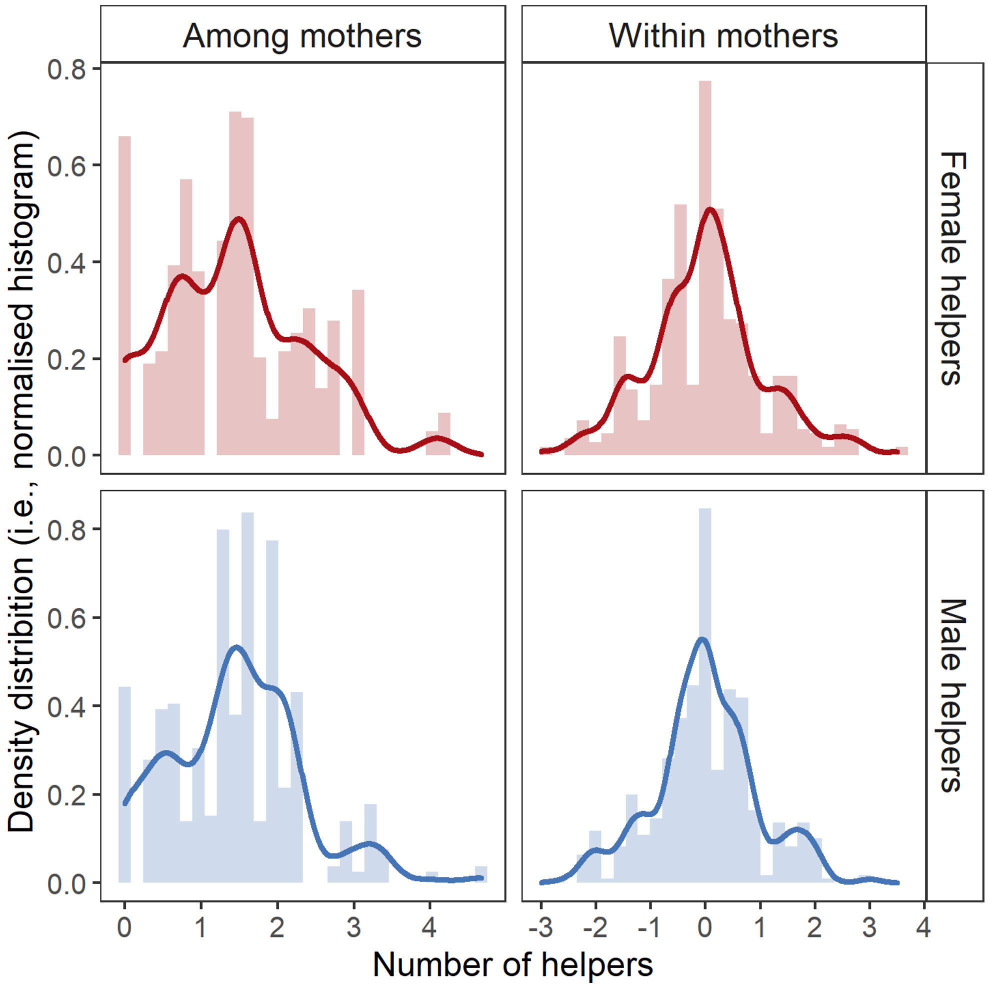

**Figure S6. Distribution of the number of female and male helpers within and among mothers** in the dataset used for egg volume analysis. Analogous distributions for female and male indicate that the power to detect female and male effects was similar.

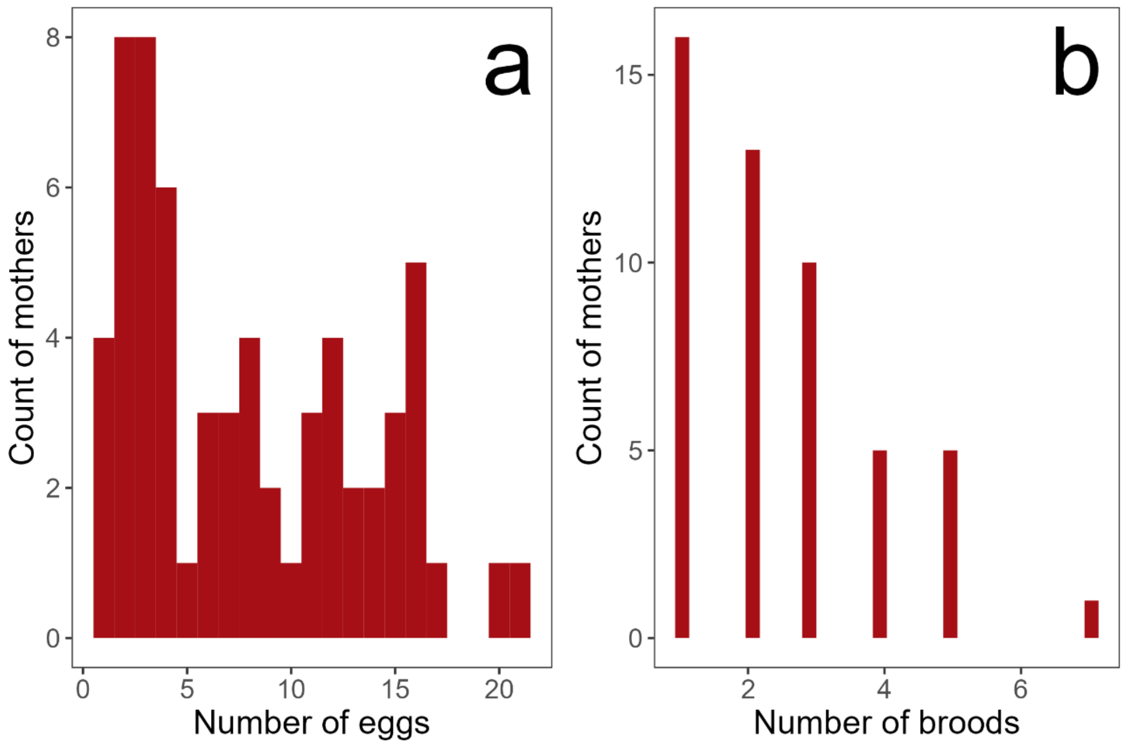

**Figure S7. Distribution of observation of egg volume and provisioning effort per female**. (a) Number of eggs per mother included in our egg volume analysis and (b) number of broods per female in our maternal provisioning analysis**.**

### Supplementary Tables

**Table S1.** Model selection table for models explaining variation in egg volume ($cm^3$). This table presents all models within ΔAIC < 6 of the top model. Model coefficients (effect sizes ± standard errors [SE]) are shown along with number of model parameters ('k'), AIC and ΔAIC. 'Heat waves' (days above 35°C), 'Clutch size' and 'Egg position' were mean centered and scaled by one standard deviation prior model fit to improve model convergence. Similarly, 'Rainfall$^1$' and 'Rainfall$^2$' were fitted as orthogonal vectors and their estimates are not back transformed in this table (i.e., units do not refer to the real data scale).

| Intercept | Number of helping females | Number of helping males | Clutch size | Egg position | Rainfall$^1$ | Rainfall$^2$ | Heat waves | Number of helping females x Clutch size | Number of helping males x Clutch size | Number of helping females x Egg position | Number of helping males x Egg position | k | AIC | ΔAIC |
|---|---|---|---|---|---|---|---|---|---|---|---|---|---|---|
| 3.64 ± 0.04 | 0.02 ± 0.01 | | | −0.04 ± 0.01 | −0.55 ± 0.22 | −0.86 ± 0.21 | −0.04 ± 0.01 | | | | | 11 | −98.49 | 0.00 |
| 3.63 ± 0.04 | 0.02 ± 0.01 | 0.01 ± 0.01 | | −0.04 ± 0.01 | −0.55 ± 0.22 | −0.86 ± 0.21 | −0.04 ± 0.01 | | | | | 12 | −97.09 | 1.40 |
| 3.64 ± 0.04 | 0.02 ± 0.01 | | | −0.05 ± 0.01 | −0.56 ± 0.22 | −0.87 ± 0.21 | −0.04 ± 0.01 | | | 0.00 ± 0.01 | | 12 | −96.89 | 1.59 |
| 3.64 ± 0.04 | 0.02 ± 0.01 | | 0.00 ± 0.01 | −0.04 ± 0.01 | −0.55 ± 0.22 | −0.86 ± 0.21 | −0.04 ± 0.01 | | | | | 12 | −96.49 | 2.00 |
| 3.63 ± 0.04 | 0.02 ± 0.01 | 0.01 ± 0.01 | | −0.05 ± 0.01 | −0.55 ± 0.22 | −0.86 ± 0.21 | −0.04 ± 0.01 | | | 0.00 ± 0.01 | | 13 | −95.46 | 3.03 |
| 3.63 ± 0.04 | 0.02 ± 0.01 | 0.01 ± 0.01 | | −0.04 ± 0.01 | −0.55 ± 0.22 | −0.86 ± 0.21 | −0.04 ± 0.01 | | | | 0.00 ± 0.01 | 13 | −95.33 | 3.16 |
| 3.63 ± 0.04 | 0.02 ± 0.01 | 0.01 ± 0.01 | 0.00 ± 0.01 | −0.04 ± 0.01 | −0.55 ± 0.22 | −0.85 ± 0.21 | −0.04 ± 0.01 | | | | | 13 | −95.10 | 3.39 |
| 3.64 ± 0.04 | 0.02 ± 0.01 | | −0.01 ± 0.01 | −0.04 ± 0.01 | −0.55 ± 0.22 | −0.87 ± 0.21 | −0.04 ± 0.01 | 0.01 ± 0.01 | | | | 13 | −95.02 | 3.47 |
| 3.64 ± 0.04 | 0.02 ± 0.01 | | 0.00 ± 0.01 | −0.05 ± 0.01 | −0.56 ± 0.22 | −0.87 ± 0.21 | −0.04 ± 0.01 | | | 0.00 ± 0.01 | | 13 | −94.89 | 3.59 |

| | | | | | | | | | | | | | | |
|---|---|---|---|---|---|---|---|---|---|---|---|---|---|---|
| 3.64 ± 0.04 | | 0.01 ± 0.01 | | −0.04 ± 0.01 | −0.54 ± 0.22 | −0.85 ± 0.22 | −0.04 ± 0.01 | | | | | 11 | −94.78 | 3.71 |
| 3.66 ± 0.04 | | | | −0.04 ± 0.01 | −0.58 ± 0.22 | −0.84 ± 0.22 | −0.04 ± 0.01 | | | | | 10 | −94.43 | 4.05 |
| 3.63 ± 0.04 | 0.02 ± 0.01 | 0.01 ± 0.01 | | −0.04 ± 0.01 | −0.55 ± 0.22 | −0.87 ± 0.21 | −0.04 ± 0.01 | | | 0.01 ± 0.01 | −0.01 ± 0.01 | 14 | −94.05 | 4.44 |
| 3.63 ± 0.04 | 0.02 ± 0.01 | 0.01 ± 0.01 | 0.00 ± 0.01 | −0.04 ± 0.01 | −0.54 ± 0.22 | −0.86 ± 0.21 | −0.04 ± 0.01 | 0.01 ± 0.01 | | | | 14 | −93.55 | 4.93 |
| 3.63 ± 0.04 | 0.02 ± 0.01 | 0.01 ± 0.01 | 0.00 ± 0.01 | −0.05 ± 0.01 | −0.55 ± 0.22 | −0.86 ± 0.21 | −0.04 ± 0.01 | | | 0.00 ± 0.01 | | 14 | −93.48 | 5.01 |
| 3.63 ± 0.04 | 0.02 ± 0.01 | 0.01 ± 0.01 | 0.01 ± 0.01 | −0.04 ± 0.01 | −0.54 ± 0.22 | −0.86 ± 0.21 | −0.04 ± 0.01 | | −0.01 ± 0.01 | | | 14 | −93.41 | 5.08 |
| 3.63 ± 0.04 | 0.02 ± 0.01 | 0.01 ± 0.01 | 0.00 ± 0.01 | −0.04 ± 0.01 | −0.55 ± 0.22 | −0.86 ± 0.21 | −0.04 ± 0.01 | | | | 0.00 ± 0.01 | 14 | −93.33 | 5.15 |
| 3.64 ± 0.04 | 0.02 ± 0.01 | | 0.00 ± 0.01 | −0.05 ± 0.01 | −0.55 ± 0.22 | −0.87 ± 0.21 | −0.04 ± 0.01 | 0.00 ± 0.01 | | 0.00 ± 0.01 | | 14 | −93.16 | 5.33 |
| 3.64 ± 0.04 | | 0.01 ± 0.01 | | −0.04 ± 0.01 | −0.54 ± 0.22 | −0.85 ± 0.22 | −0.04 ± 0.01 | | | | 0.00 ± 0.01 | 12 | −92.97 | 5.51 |
| 3.64 ± 0.04 | | 0.01 ± 0.01 | 0.00 ± 0.01 | −0.04 ± 0.01 | −0.54 ± 0.22 | −0.85 ± 0.22 | −0.04 ± 0.01 | | | | | 12 | −92.79 | 5.70 |
| 3.66 ± 0.04 | | | 0.00 ± 0.01 | −0.04 ± 0.01 | −0.57 ± 0.22 | −0.84 ± 0.22 | −0.04 ± 0.01 | | | | | 11 | −92.52 | 5.97 |

6

**Table S2.** Model selection table for models explaining variation in egg volume (cm$^3$), when population-level variation in female and male helper number were partitioned into their within-mother (Δ) and among-mother (μ) components prior to model selection. The table presents all models within ΔAIC < 6 of the top model. Model coefficients (effect sizes ± standard errors [SE]) are shown along with number of model parameters (‘k’), AIC and ΔAIC. ‘Heat waves’ (days above 35˚C), ‘Clutch size’ and ‘Egg position’ were mean centered and scaled by one standard deviation prior model fit to improve model convergence. Similarly, ‘Rainfall$^1$’ and ‘Rainfall$^2$’ were fitted as orthogonal vectors and their estimates are not back transformed in this table (i.e., units do not refer to the real data scale).

| Intercept | Δ Number of helping females | μ Number of helping females | Δ Number of helping males | μ Number of helping males | Clutch size | Egg position | Rainfall$^1$ | Rainfall$^2$ | Heat waves | k | AIC | ΔAIC |
|---|---|---|---|---|---|---|---|---|---|---|---|---|
| 3.66 ± 0.04 | 0.02 ± 0.01 | | | | | −0.04 ± 0.01 | −0.55 ± 0.22 | −0.87 ± 0.21 | −0.04 ± 0.01 | 11 | −98.33 | 0.00 |
| 3.66 ± 0.04 | 0.02 ± 0.01 | | 0.01 ± 0.01 | | | −0.04 ± 0.01 | −0.54 ± 0.22 | −0.87 ± 0.21 | −0.04 ± 0.01 | 12 | −96.88 | 1.45 |
| 3.65 ± 0.05 | 0.02 ± 0.01 | 0.01 ± 0.02 | | | | −0.04 ± 0.01 | −0.55 ± 0.22 | −0.86 ± 0.21 | −0.04 ± 0.01 | 12 | −96.61 | 1.72 |
| 3.65 ± 0.05 | 0.02 ± 0.01 | | | 0.01 ± 0.02 | | −0.04 ± 0.01 | −0.55 ± 0.22 | −0.87 ± 0.21 | −0.04 ± 0.01 | 12 | −96.59 | 1.74 |
| 3.66 ± 0.04 | 0.02 ± 0.01 | | | | 0.00 ± 0.01 | −0.04 ± 0.01 | −0.55 ± 0.22 | −0.87 ± 0.21 | −0.04 ± 0.01 | 12 | −96.33 | 2.00 |
| 3.65 ± 0.05 | 0.02 ± 0.01 | 0.01 ± 0.02 | 0.01 ± 0.01 | | | −0.04 ± 0.01 | −0.54 ± 0.22 | −0.86 ± 0.21 | −0.04 ± 0.01 | 13 | −95.17 | 3.16 |
| 3.65 ± 0.05 | 0.02 ± 0.01 | | 0.01 ± 0.01 | 0.01 ± 0.02 | | −0.04 ± 0.01 | −0.54 ± 0.22 | −0.86 ± 0.21 | −0.04 ± 0.01 | 13 | −95.15 | 3.18 |
| 3.66 ± 0.04 | 0.02 ± 0.01 | | 0.01 ± 0.01 | | 0.00 ± 0.01 | −0.04 ± 0.01 | −0.54 ± 0.22 | −0.87 ± 0.21 | −0.04 ± 0.01 | 13 | −94.88 | 3.45 |
| 3.64 ± 0.05 | 0.02 ± 0.01 | 0.01 ± 0.03 | | 0.01 ± 0.03 | | −0.04 ± 0.01 | −0.55 ± 0.22 | −0.86 ± 0.21 | −0.04 ± 0.01 | 13 | −94.69 | 3.64 |

| | | | | | | | | | | | | |
|---|---|---|---|---|---|---|---|---|---|---|---|---|
| 3.65 ± 0.05 | 0.02 ± 0.01 | 0.01 ± 0.02 | | | 0.00 ± 0.01 | −0.04 ± 0.01 | −0.55 ± 0.22 | −0.86 ± 0.21 | −0.04 ± 0.01 | 13 | −94.61 | 3.72 |
| 3.65 ± 0.05 | 0.02 ± 0.01 | | | 0.01 ± 0.02 | 0.00 ± 0.01 | −0.04 ± 0.01 | −0.55 ± 0.22 | −0.87 ± 0.21 | −0.04 ± 0.01 | 13 | −94.59 | 3.74 |
| 3.66 ± 0.04 | | | 0.01 ± 0.01 | | | −0.04 ± 0.01 | −0.54 ± 0.22 | −0.85 ± 0.22 | −0.04 ± 0.01 | 11 | −94.54 | 3.79 |
| 3.66 ± 0.04 | | | | | | −0.04 ± 0.01 | −0.58 ± 0.22 | −0.84 ± 0.22 | −0.04 ± 0.01 | 10 | −94.43 | 3.90 |
| 3.64 ± 0.05 | 0.02 ± 0.01 | 0.01 ± 0.03 | 0.01 ± 0.01 | 0.01 ± 0.03 | | −0.04 ± 0.01 | −0.54 ± 0.22 | −0.86 ± 0.21 | −0.04 ± 0.01 | 14 | −93.25 | 5.08 |
| 3.65 ± 0.05 | 0.02 ± 0.01 | 0.01 ± 0.02 | 0.01 ± 0.01 | | 0.00 ± 0.01 | −0.04 ± 0.01 | −0.54 ± 0.22 | −0.86 ± 0.21 | −0.04 ± 0.01 | 14 | −93.17 | 5.16 |
| 3.65 ± 0.05 | 0.02 ± 0.01 | | 0.01 ± 0.01 | 0.01 ± 0.02 | 0.00 ± 0.01 | −0.04 ± 0.01 | −0.54 ± 0.22 | −0.86 ± 0.21 | −0.04 ± 0.01 | 14 | −93.16 | 5.17 |
| 3.65 ± 0.05 | | 0.01 ± 0.02 | 0.01 ± 0.01 | | | −0.04 ± 0.01 | −0.54 ± 0.22 | −0.85 ± 0.22 | −0.04 ± 0.01 | 12 | −92.81 | 5.52 |
| 3.65 ± 0.05 | | | 0.01 ± 0.01 | 0.01 ± 0.02 | | −0.04 ± 0.01 | −0.54 ± 0.22 | −0.85 ± 0.22 | −0.04 ± 0.01 | 12 | −92.79 | 5.54 |
| 3.64 ± 0.05 | 0.02 ± 0.01 | 0.01 ± 0.03 | | 0.01 ± 0.03 | 0.00 ± 0.01 | −0.04 ± 0.01 | −0.55 ± 0.22 | −0.86 ± 0.21 | −0.04 ± 0.01 | 14 | −92.69 | 5.64 |
| 3.65 ± 0.05 | | 0.01 ± 0.02 | | | | −0.04 ± 0.01 | −0.57 ± 0.22 | −0.84 ± 0.22 | −0.04 ± 0.01 | 11 | −92.59 | 5.74 |
| 3.66 ± 0.04 | | | 0.01 ± 0.01 | | 0.00 ± 0.01 | −0.04 ± 0.01 | −0.53 ± 0.22 | −0.85 ± 0.22 | −0.04 ± 0.01 | 12 | −92.57 | 5.76 |
| 3.65 ± 0.05 | | | | 0.01 ± 0.02 | | −0.04 ± 0.01 | −0.57 ± 0.22 | −0.84 ± 0.22 | −0.04 ± 0.01 | 11 | −92.56 | 5.77 |

| | | | | | | | | |
|---|---|---|---|---|---|---|---|---|
| 3.66 ± 0.04 | 0.00 ± 0.01 | −0.04 ± 0.01 | −0.57 ± 0.22 | −0.84 ± 0.22 | −0.04 ± 0.01 | 11 | −92.52 | 5.81 |

**Table S3.** Model selection table for models explaining variation in maternal provisioning rate (feeds / hour). This table presents all models within ΔAIC < 6 of the top model. Model coefficients (effect sizes ± standard errors [SE]) are shown along with number of model parameters ('k'), AIC and ΔAIC. 'Heat waves' (days above 35˚C) and 'Brood size' were mean centered and scaled by one standard deviation prior model fit to improve model convergence. Similarly, 'Rainfall$^1$' and 'Rainfall$^2$' were fitted as orthogonal vectors and their estimates are not back transformed in this table (i.e., units do not refer to the real data scale).

| Intercept | Number of helping females | Number of helping males | Brood size | Rainfall$^1$ | Rainfall$^2$ | Heat waves | Number of helping females x Brood size | Number of helping males x Brood size | k | AIC | ΔAIC |
|---|---|---|---|---|---|---|---|---|---|---|---|
| 7.37 ± 0.47 | −0.46 ± 0.19 | | 1.45 ± 0.24 | 4.60 ± 3.02 | 7.69 ± 2.75 | 0.71 ± 0.30 | | | 10 | 606.99 | 0.00 |
| 7.46 ± 0.54 | −0.46 ± 0.19 | −0.07 ± 0.23 | 1.44 ± 0.24 | 4.55 ± 3.02 | 7.67 ± 2.75 | 0.70 ± 0.30 | | | 11 | 608.89 | 1.90 |
| 7.37 ± 0.47 | −0.46 ± 0.20 | | 1.43 ± 0.32 | 4.64 ± 3.09 | 7.68 ± 2.76 | 0.71 ± 0.30 | 0.01 ± 0.20 | | 11 | 608.99 | 2.00 |
| 7.46 ± 0.51 | −0.44 ± 0.19 | −0.13 ± 0.23 | 1.81 ± 0.36 | 4.45 ± 2.96 | 7.85 ± 2.73 | 0.71 ± 0.29 | | −0.30 ± 0.20 | 12 | 609.01 | 2.02 |
| 7.35 ± 0.52 | −0.47 ± 0.20 | | 1.47 ± 0.25 | 7.59 ± 2.79 | 8.79 ± 2.77 | | | | 9 | 610.26 | 3.27 |
| 6.82 ± 0.43 | | | 1.47 ± 0.25 | 4.66 ± 3.10 | 7.64 ± 2.81 | 0.71 ± 0.31 | | | 9 | 610.47 | 3.48 |
| 7.46 ± 0.51 | −0.43 ± 0.20 | −0.13 ± 0.23 | 1.75 ± 0.39 | 4.75 ± 3.03 | 7.78 ± 2.73 | 0.69 ± 0.29 | 0.10 ± 0.20 | −0.32 ± 0.21 | 13 | 610.79 | 3.80 |
| 7.46 ± 0.54 | −0.45 ± 0.20 | −0.07 ± 0.23 | 1.42 ± 0.33 | 4.59 ± 3.09 | 7.66 ± 2.75 | 0.70 ± 0.30 | 0.01 ± 0.20 | | 12 | 610.89 | 3.90 |
| 7.52 ± 0.58 | −0.46 ± 0.20 | −0.14 ± 0.23 | 1.45 ± 0.25 | 7.38 ± 2.80 | 8.72 ± 2.77 | | | | 10 | 611.92 | 4.93 |
| 7.01 ± 0.48 | | −0.19 ± 0.23 | 1.86 ± 0.36 | 4.59 ± 3.02 | 7.82 ± 2.78 | 0.67 ± 0.30 | | −0.32 ± 0.21 | 11 | 612.13 | 5.14 |

| | | | | | | | | | | | |
|---|---|---|---|---|---|---|---|---|---|---|---|
| 7.35 ± 0.52 | −0.46 ± 0.20 | | 1.41 ± 0.33 | 7.73 ± 2.84 | 8.73 ± 2.79 | | 0.05 ± 0.20 | | 10 | 612.20 | 5.21 |
| 6.96 ± 0.51 | | −0.11 ± 0.23 | 1.46 ± 0.25 | 4.62 ± 3.09 | 7.59 ± 2.81 | 0.70 ± 0.31 | | | 10 | 612.29 | 5.30 |
| 7.55 ± 0.56 | −0.44 ± 0.20 | −0.20 ± 0.24 | 1.78 ± 0.36 | 7.16 ± 2.76 | 8.91 ± 2.75 | | | −0.28 ± 0.21 | 11 | 612.30 | 5.31 |
| 7.39 ± 0.44 | −0.45 ± 0.20 | | 1.30 ± 0.24 | 4.82 ± 3.06 | | 0.82 ± 0.30 | | | 9 | 612.35 | 5.36 |
| 7.43 ± 0.47 | −0.46 ± 0.20 | | 1.33 ± 0.24 | | | 1.02 ± 0.28 | | | 8 | 612.67 | 5.68 |

**Table S4.** Model selection table for models explaining variation in maternal provisioning rate (feeds / hour), when population-level variation in female and male helper number were partitioned into their within-mother (Δ) and among-mother (μ) components prior to model selection. Model coefficients (effect sizes ± standard errors) are shown along with number of model parameters (‘k’), AIC and ΔAIC. ‘Heat waves’ (days above 35˚C) and ‘Brood size’ were mean centered and scaled by one standard deviation prior model fit to improve model convergence. Similarly, ‘Rainfall$^1$’ and ‘Rainfall$^2$’ were fitted as orthogonal vectors and their estimates are not back transformed in this table (i.e., units do not refer to the real data scale).

| Intercept | Δ Number of helping females | μ Number of helping females | Δ Number of helping males | μ Number of helping males | Brood size | Rainfall$^1$ | Rainfall$^2$ | Heat waves | k | AIC | ΔAIC |
|---|---|---|---|---|---|---|---|---|---|---|---|
| 6.85 ± 0.41 | −0.53 ± 0.26 | | | | 1.48 ± 0.24 | 4.62 ± 3.04 | 7.74 ± 2.76 | 0.73 ± 0.30 | 10 | 608.40 | 0.00 |
| 7.27 ± 0.52 | −0.54 ± 0.26 | −0.36 ± 0.28 | | | 1.45 ± 0.24 | 4.59 ± 3.02 | 7.72 ± 2.75 | 0.72 ± 0.30 | 11 | 608.78 | 0.37 |
| 7.24 ± 0.61 | −0.56 ± 0.26 | | | −0.32 ± 0.39 | 1.46 ± 0.24 | 4.61 ± 3.02 | 7.78 ± 2.76 | 0.72 ± 0.30 | 11 | 609.77 | 1.36 |
| 6.85 ± 0.41 | −0.55 ± 0.27 | | 0.06 ± 0.30 | | 1.48 ± 0.24 | 4.66 ± 3.04 | 7.76 ± 2.77 | 0.74 ± 0.30 | 11 | 610.36 | 1.96 |
| 6.82 ± 0.43 | | | | | 1.47 ± 0.25 | 4.66 ± 3.10 | 7.64 ± 2.81 | 0.71 ± 0.31 | 9 | 610.47 | 2.07 |
| 7.45 ± 0.64 | −0.56 ± 0.26 | −0.33 ± 0.30 | | −0.19 ± 0.40 | 1.45 ± 0.24 | 4.59 ± 3.01 | 7.73 ± 2.74 | 0.71 ± 0.30 | 12 | 610.56 | 2.16 |
| 7.27 ± 0.53 | −0.54 ± 0.27 | −0.37 ± 0.29 | −0.01 ± 0.30 | | 1.45 ± 0.24 | 4.58 ± 3.02 | 7.71 ± 2.75 | 0.72 ± 0.30 | 12 | 610.77 | 2.37 |
| 7.22 ± 0.53 | | −0.35 ± 0.29 | | | 1.45 ± 0.25 | 4.66 ± 3.07 | 7.60 ± 2.79 | 0.70 ± 0.31 | 10 | 611.01 | 2.61 |
| 7.23 ± 0.61 | −0.58 ± 0.27 | | 0.08 ± 0.30 | −0.32 ± 0.39 | 1.46 ± 0.24 | 4.67 ± 3.03 | 7.80 ± 2.76 | 0.73 ± 0.30 | 12 | 611.70 | 3.30 |

| | | | | | | | | | | | |
|---|---|---|---|---|---|---|---|---|---|---|---|
| 6.83 ± 0.46 | −0.51 ± 0.27 | | | | 1.49 ± 0.25 | 7.67 ± 2.80 | 8.83 ± 2.79 | | 9 | 611.98 | 3.58 |
| 7.27 ± 0.57 | −0.53 ± 0.27 | −0.40 ± 0.29 | | | 1.47 ± 0.25 | 7.61 ± 2.79 | 8.82 ± 2.78 | | 10 | 612.15 | 3.74 |
| 7.05 ± 0.62 | | | | −0.20 ± 0.40 | 1.46 ± 0.25 | 4.69 ± 3.09 | 7.64 ± 2.81 | 0.70 ± 0.31 | 10 | 612.24 | 3.84 |
| 6.83 ± 0.42 | | | −0.06 ± 0.30 | | 1.47 ± 0.25 | 4.63 ± 3.10 | 7.61 ± 2.81 | 0.71 ± 0.31 | 10 | 612.44 | 4.03 |
| 7.44 ± 0.64 | −0.56 ± 0.27 | −0.33 ± 0.30 | 0.01 ± 0.31 | −0.19 ± 0.41 | 1.45 ± 0.24 | 4.60 ± 3.02 | 7.74 ± 2.75 | 0.71 ± 0.30 | 13 | 612.56 | 4.16 |
| 7.26 ± 0.54 | | −0.38 ± 0.29 | −0.14 ± 0.30 | | 1.44 ± 0.25 | 4.58 ± 3.08 | 7.54 ± 2.79 | 0.68 ± 0.31 | 11 | 612.81 | 4.41 |
| 7.28 ± 0.65 | | −0.34 ± 0.30 | | −0.06 ± 0.41 | 1.44 ± 0.25 | 4.67 ± 3.07 | 7.60 ± 2.79 | 0.70 ± 0.31 | 11 | 612.99 | 4.58 |
| 7.29 ± 0.66 | −0.55 ± 0.27 | | | −0.38 ± 0.41 | 1.47 ± 0.25 | 7.59 ± 2.79 | 8.87 ± 2.78 | | 10 | 613.13 | 4.72 |
| 6.81 ± 0.46 | | | | | 1.48 ± 0.25 | 7.68 ± 2.84 | 8.66 ± 2.83 | | 8 | 613.63 | 5.22 |
| 6.87 ± 0.38 | −0.51 ± 0.27 | | | | 1.33 ± 0.25 | 4.83 ± 3.09 | | 0.84 ± 0.30 | 9 | 613.85 | 5.44 |
| 7.49 ± 0.69 | −0.55 ± 0.27 | −0.35 ± 0.30 | | −0.23 ± 0.42 | 1.46 ± 0.25 | 7.57 ± 2.78 | 8.83 ± 2.77 | | 11 | 613.86 | 5.45 |
| 6.84 ± 0.46 | −0.51 ± 0.27 | | −0.03 ± 0.30 | | 1.49 ± 0.25 | 7.63 ± 2.82 | 8.81 ± 2.79 | | 10 | 613.97 | 5.56 |
| 7.23 ± 0.57 | | −0.38 ± 0.30 | | | 1.46 ± 0.25 | 7.61 ± 2.83 | 8.62 ± 2.82 | | 9 | 614.01 | 5.61 |

| | | | | | | | | | | | |
|---|---|---|---|---|---|---|---|---|---|---|---|
| 7.31 ± 0.58 | −0.51 ± 0.27 | −0.42 ± 0.30 | −0.11 ± 0.31 | | 1.46 ± 0.25 | 7.47 ± 2.81 | 8.76 ± 2.78 | | 11 | 614.03 | 5.62 |
| 6.91 ± 0.42 | −0.52 ± 0.27 | | | | 1.36 ± 0.25 | | | 1.04 ± 0.28 | 8 | 614.11 | 5.70 |
| 7.06 ± 0.62 | | | −0.06 ± 0.30 | −0.20 ± 0.40 | 1.45 ± 0.25 | 4.66 ± 3.09 | 7.62 ± 2.81 | 0.69 ± 0.31 | 11 | 614.21 | 5.80 |
| 7.31 ± 0.50 | −0.52 ± 0.27 | −0.38 ± 0.29 | | | 1.30 ± 0.24 | 4.80 ± 3.06 | | 0.83 ± 0.30 | 10 | 614.23 | 5.82 |

**Table S5.** Model selection table for models explaining variation in clutch size (zero-truncated models). This analysis revealed very week evidence for an effect of female or male helper number on clutch size. Models including female or male helper number received similar support from the data that the intercept-only model. Model coefficients (effect sizes ± standard errors) are shown along with number of model parameters (‘k’), AIC and ΔAIC. ‘Clutch order’ was mean centered and scaled by one standard deviation prior model fit to improve model convergence.

| Intercept | Number of helping females | Number of helping males | Clutch order | k | AIC | ΔAIC |
|---|---|---|---|---|---|---|
| 0.63 ± 0.06 | −0.06 ± 0.04 | | | 5 | 851.40 | 0.00 |
| 0.63 ± 0.07 | | −0.07 ± 0.04 | | 5 | 851.86 | 0.46 |
| 0.55 ± 0.05 | | | | 4 | 851.95 | 0.55 |
| 0.66 ± 0.07 | −0.05 ± 0.04 | −0.05 ± 0.05 | | 6 | 852.40 | 1.00 |
| 0.62 ± 0.06 | −0.06 ± 0.04 | | 0.04 ± 0.05 | 6 | 852.74 | 1.34 |
| 0.55 ± 0.05 | | | 0.04 ± 0.05 | 5 | 853.08 | 1.68 |
| 0.62 ± 0.07 | | −0.06 ± 0.04 | 0.04 ± 0.05 | 6 | 853.18 | 1.78 |
| 0.65 ± 0.07 | −0.05 ± 0.04 | −0.04 ± 0.05 | 0.03 ± 0.05 | 7 | 853.89 | 2.49 |

**Table S6**. Model selection table for models explaining variation in clutch size (zero-truncated models) after partitioning variation in female and male helper number into their within-mother (Δ) and among-mother (μ) components. Models including female or male helper number components received similar support from the data that the intercept-only model. Model coefficients (effect sizes ± standard errors) are shown along with number of model parameters (‘k’), AIC and ΔAIC. ‘Clutch order’ was mean centered and scaled by one standard deviation prior model fit to improve model convergence.

| Intercept | Δ Number of helping females | μ Number of helping females | Δ Number of helping males | μ Number of helping males | Clutch order | k | AIC | ΔAIC |
|---|---|---|---|---|---|---|---|---|
| 0.55 ± 0.05 | −0.07 ± 0.05 | | | | | 5 | 851.94 | 0.00 |
| 0.55 ± 0.05 | | | | | | 4 | 851.95 | 0.01 |
| 0.55 ± 0.05 | | | −0.08 ± 0.05 | | | 5 | 852.07 | 0.13 |
| 0.55 ± 0.05 | | | | | 0.04 ± 0.05 | 5 | 853.08 | 1.14 |
| 0.55 ± 0.05 | −0.05 ± 0.05 | | −0.05 ± 0.06 | | | 6 | 853.11 | 1.17 |
| 0.66 ± 0.14 | −0.07 ± 0.05 | −0.05 ± 0.06 | | | | 6 | 853.32 | 1.38 |
| 0.55 ± 0.05 | −0.06 ± 0.05 | | | | 0.04 ± 0.05 | 6 | 853.35 | 1.41 |
| 0.66 ± 0.14 | | −0.05 ± 0.06 | −0.08 ± 0.05 | | | 6 | 853.49 | 1.55 |
| 0.55 ± 0.05 | | | −0.07 ± 0.05 | | 0.03 ± 0.05 | 6 | 853.52 | 1.58 |
| 0.64 ± 0.15 | | −0.04 ± 0.07 | | | | 5 | 853.57 | 1.63 |
| 0.64 ± 0.16 | −0.07 ± 0.05 | | | −0.04 ± 0.07 | | 6 | 853.59 | 1.65 |
| 0.65 ± 0.16 | | | −0.08 ± 0.05 | −0.04 ± 0.07 | | 6 | 853.69 | 1.75 |

| | | | | | | | | |
|---|---|---|---|---|---|---|---|---|
| 0.63 ± 0.16 | | | | −0.04 ± 0.07 | | 5 | 853.71 | 1.77 |
| 0.66 ± 0.14 | −0.05 ± 0.05 | −0.05 ± 0.06 | −0.05 ± 0.06 | | | 7 | 854.48 | 2.54 |
| 0.65 ± 0.15 | | −0.05 ± 0.07 | | | 0.05 ± 0.05 | 6 | 854.59 | 2.65 |
| 0.55 ± 0.05 | −0.05 ± 0.05 | | −0.05 ± 0.06 | | 0.03 ± 0.05 | 7 | 854.70 | 2.76 |
| 0.66 ± 0.14 | −0.06 ± 0.05 | −0.05 ± 0.06 | | | 0.04 ± 0.05 | 7 | 854.71 | 2.77 |
| 0.64 ± 0.16 | −0.05 ± 0.05 | | −0.05 ± 0.06 | −0.04 ± 0.07 | | 7 | 854.73 | 2.79 |
| 0.64 ± 0.16 | | | | −0.04 ± 0.07 | 0.05 ± 0.05 | 6 | 854.75 | 2.81 |
| 0.66 ± 0.14 | | −0.05 ± 0.06 | −0.07 ± 0.05 | | 0.04 ± 0.05 | 7 | 854.92 | 2.98 |
| 0.65 ± 0.16 | −0.06 ± 0.05 | | | −0.04 ± 0.07 | 0.04 ± 0.05 | 7 | 854.95 | 3.01 |
| 0.65 ± 0.16 | | | −0.07 ± 0.05 | −0.05 ± 0.07 | 0.04 ± 0.05 | 7 | 855.10 | 3.16 |
| 0.71 ± 0.19 | −0.07 ± 0.05 | −0.04 ± 0.06 | | −0.03 ± 0.07 | | 7 | 855.15 | 3.21 |
| 0.71 ± 0.19 | | −0.04 ± 0.06 | −0.08 ± 0.05 | −0.03 ± 0.07 | | 7 | 855.30 | 3.36 |
| 0.69 ± 0.20 | | −0.04 ± 0.07 | | −0.03 ± 0.07 | | 6 | 855.43 | 3.49 |
| 0.66 ± 0.14 | −0.05 ± 0.05 | −0.05 ± 0.06 | −0.05 ± 0.06 | | 0.03 ± 0.05 | 8 | 856.06 | 4.12 |

| | | | | | | | | |
|---|---|---|---|---|---|---|---|---|
| 0.65 ± 0.16 | −0.05 ± 0.05 | | −0.05 ± 0.06 | −0.05 ± 0.07 | 0.03 ± 0.05 | 8 | 856.29 | 4.35 |
| 0.71 ± 0.19 | −0.05 ± 0.05 | −0.04 ± 0.06 | −0.05 ± 0.06 | −0.03 ± 0.07 | | 8 | 856.30 | 4.36 |
| 0.71 ± 0.20 | | −0.04 ± 0.07 | | −0.03 ± 0.07 | 0.05 ± 0.05 | 7 | 856.40 | 4.46 |
| 0.71 ± 0.19 | −0.06 ± 0.05 | −0.04 ± 0.06 | | −0.03 ± 0.07 | 0.04 ± 0.05 | 8 | 856.52 | 4.58 |
| 0.71 ± 0.19 | | −0.04 ± 0.06 | −0.07 ± 0.05 | −0.03 ± 0.07 | 0.04 ± 0.05 | 8 | 856.70 | 4.76 |
| 0.71 ± 0.19 | −0.05 ± 0.05 | −0.04 ± 0.06 | −0.05 ± 0.06 | −0.03 ± 0.07 | 0.03 ± 0.05 | 9 | 857.86 | 5.92 |

**Table S7.** Model selection table for models explaining variation in the number of clutches laid per year. The model with 'Rainfall' as a single predictor received the strongest support from the data. Removing this term caused a decrease in AIC of 10.12. Model coefficients (effect sizes ± standard errors) are shown along with number of model parameters ('k'), AIC and ΔAIC. Estimates for 'Rainfall' are given for 100 mm of rainfall (e.g., change in number of clutches per 100mm of rainfall).

| Intercept | Number of helping females | Number of helping males | Rainfall | k | AIC | ΔAIC |
|---|---|---|---|---|---|---|
| 0.48 ± 0.12 | | | 0.11 ± 0.03 | 4 | 710.45 | 0.00 |
| 0.43 ± 0.13 | | 0.04 ± 0.04 | 0.11 ± 0.03 | 5 | 711.48 | 1.03 |
| 0.47 ± 0.12 | 0.01 ± 0.04 | | 0.11 ± 0.03 | 5 | 712.34 | 1.89 |
| 0.43 ± 0.13 | 0.00 ± 0.04 | 0.04 ± 0.05 | 0.11 ± 0.03 | 6 | 713.48 | 3.03 |

**Table S8**. Model selection table for models explaining variation in the number of clutches laid per year after partitioning variation in female and male helper number into their within-mother (Δ) and among-mother (μ) components. Model coefficients (effect sizes ± standard errors) are shown along with number of model parameters (‘k’), AIC and ΔAIC. Estimates for ‘Rainfall’ are given for 100mm of rainfall (e.g., change in number of clutches per 100mm of rainfall).

| Intercept | Δ Number of helping females | μ Number of helping females | Δ Number of helping males | μ Number of helping males | Rainfall | k | AIC | ΔAIC |
|---|---|---|---|---|---|---|---|---|
| −0.05 ± 0.22 | | | | 0.24 ± 0.08 | 0.11 ± 0.03 | 5 | 704.16 | 0.00 |
| −0.08 ± 0.22 | −0.06 ± 0.05 | | | 0.25 ± 0.08 | 0.12 ± 0.03 | 6 | 704.70 | 0.54 |
| −0.13 ± 0.23 | | 0.08 ± 0.07 | | 0.20 ± 0.09 | 0.11 ± 0.03 | 6 | 704.94 | 0.78 |
| −0.06 ± 0.22 | | | −0.05 ± 0.05 | 0.24 ± 0.08 | 0.11 ± 0.03 | 6 | 705.14 | 0.98 |
| −0.18 ± 0.24 | −0.06 ± 0.05 | 0.09 ± 0.07 | | 0.21 ± 0.09 | 0.11 ± 0.03 | 7 | 705.17 | 1.02 |
| −0.14 ± 0.23 | | 0.08 ± 0.07 | −0.05 ± 0.05 | 0.21 ± 0.09 | 0.11 ± 0.03 | 7 | 705.93 | 1.77 |
| −0.08 ± 0.22 | −0.05 ± 0.05 | | −0.04 ± 0.06 | 0.25 ± 0.08 | 0.12 ± 0.03 | 7 | 706.14 | 1.98 |
| −0.18 ± 0.24 | −0.06 ± 0.05 | 0.09 ± 0.07 | −0.04 ± 0.06 | 0.21 ± 0.09 | 0.11 ± 0.03 | 8 | 706.67 | 2.51 |
| 0.20 ± 0.18 | | 0.14 ± 0.07 | | | 0.10 ± 0.03 | 5 | 708.28 | 4.12 |
| 0.17 ± 0.19 | −0.06 ± 0.05 | 0.15 ± 0.07 | | | 0.11 ± 0.03 | 6 | 708.82 | 4.66 |
| 0.20 ± 0.18 | | 0.14 ± 0.07 | −0.05 ± 0.06 | | 0.10 ± 0.03 | 6 | 709.58 | 5.43 |

**Supplementary materials references**
